# Face photo-based age acceleration predicts all-cause mortality and differs among occupations

**DOI:** 10.1101/2025.04.16.649078

**Authors:** Bence Király, Iván Fejes, Csaba Kerepesi

## Abstract

While scientists argue what aging is and what drives aging, it is widely accepted that our face changes drastically with age and that mortality increases in late life. We hypothesize that people of the same age can be biologically older than others and that the human face may reflect accelerated molecular aging. To test this hypothesis we examine the associations of face photo-based age acceleration with mortality and lifestyle. For this purpose, we trained and tested artificial intelligence models on 442,110 photos of famous people. We found that face photo-based age predicts all-cause mortality for middle-aged and older individuals meaning that those age faster based on their face photo die sooner. We also found that, based on face photos, sport is the slowest aging occupation among famous people consistently to previous findings showing the benefits of exercise to epigenetic aging. Overall, we demonstrate that the face photo-base age model approaches biological age in some extent and provides a low-cost and fast complementary measurement for personalized medicine, as well as aging and rejuvenation studies. The model is available for demonstration and academic research purposes at https://photoage.sztaki.hu/.

## INTRODUCTION

Aging is a highly complex biological process that has a major impact on health, the economy, and society and is associated with numerous diseases and mortality. It would be beneficial to slow down, stop, or reverse aging through lifestyle considerations and medical interventions to lead to a longer and healthier life^1,2^. Despite the recent progress of aging research, there are still a disagreement about what aging is and what the driving processes of aging are^3^. However, it is clear and widely accepted that our face changes drastically with age coupled with increased mortality in late life. In this study, we focused on these important aspects of human aging seeking potential associations.

One of the most important directions of aging research is searching for biomarkers that are sensitive to the progress of aging and thus able to measure biological age. The practical use of biological age has the potential to help physicians in personalized medicine decisions and the evaluation of potential anti-aging treatments. Epigenetic clocks, the state-of-the-art measurements of biological age, can estimate chronological age with high accuracy from DNA methylation levels using machine learning methods^4,5^. It was shown that the age-adjusted age prediction of epigenetic clocks (so-called epigenetic age acceleration) is predictive of all-cause mortality, coronary heart diseases, and cancer^6–8^. In addition, accelerated aging has been observed in the study population for many age-related diseases^9^. These results suggest that epigenetic clocks can approach biological age more accurately than chronological age.

One of the major drawbacks of the current measurements of biological age is that they are expensive and not immediately available. It takes several days and expensive instrumentation to measure DNA methylation levels in cells. Additionally, it is a challenge to interpret epigenetic clocks. In contrast, a digital photo is easy to produce and available immediately. Here, we hypothesized that the human face reflects accelerated molecular aging. The recent advances in artificial intelligence, especially transfer learning enabled age predictions based on (2D, and 3D) facial images with accuracies comparable to epigenetic clocks^10–15^. However, the applicability of AI-predicted face age as a measurement of biological age has not been completely clear as its association with all-cause mortality has been not tested. To fill this gap, here, we use the IMDB-WIKI database that contains thousands of age-tagged photos of famous people and collected additional metadata including death dates^16^. Then we trained deep neural networks via transfer learning to predict age from face photos and tested whether age acceleration predicts all-cause mortality. As epigenetic studies found that lifestyle affects epigenetic age acceleration^2^, we also investigated that photo-based age accelerations associated with different occupations.

## RESULTS

### Development of a face photo-based age prediction model using automatically collected data of famous people

To test whether the face photo-based age acceleration is associated with mortality and lifestyle attributes, we trained and tested artificial intelligence models on 442,110 photo images of famous people available in the IMDB-WIKI database (Fig. 1a). We used 50% of the images for model development and validation, while we used the remaining 50% for exploratory analysis. The age prediction performance of the final model was r = 0.62 and MAE = 7.38 years in the test set, as well as r = 0.83 and MAE = 6.48 years in an external dataset, FG-NET, which contains mugshots of average people (Table 1, Fig. 1b).

**Table 1.**
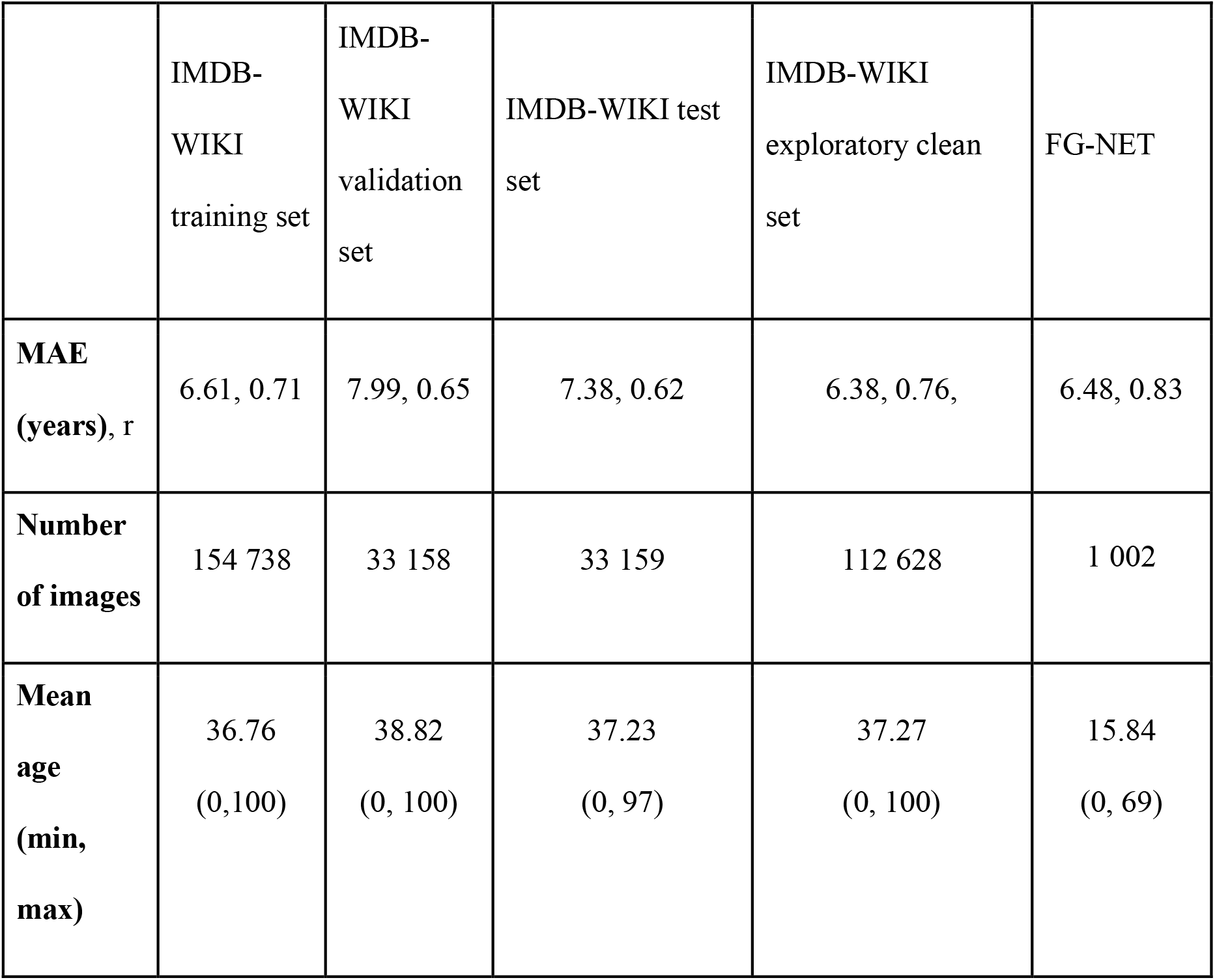
Evaluation of the face photo-based age prediction model on different datasets

|  | IMDB-WIKI training set | IMDB-WIKI validation set | IMDB-WIKI test set | IMDB-WIKI exploratory clean set | FG-NET |
| --- | --- | --- | --- | --- | --- |
| **MAE (years), r** | 6.61, 0.71 | 7.99, 0.65 | 7.38, 0.62 | 6.38, 0.76, | 6.48, 0.83 |
| **Number of images** | 154 738 | 33 158 | 33 159 | 112 628 | 1 002 |
| **Mean age (min, max)** | 36.76<br>(0,100) | 38.82<br>(0, 100) | 37.23<br>(0, 97) | 37.27<br>(0, 100) | 15.84<br>(0, 69) |

**Fig. 1.**
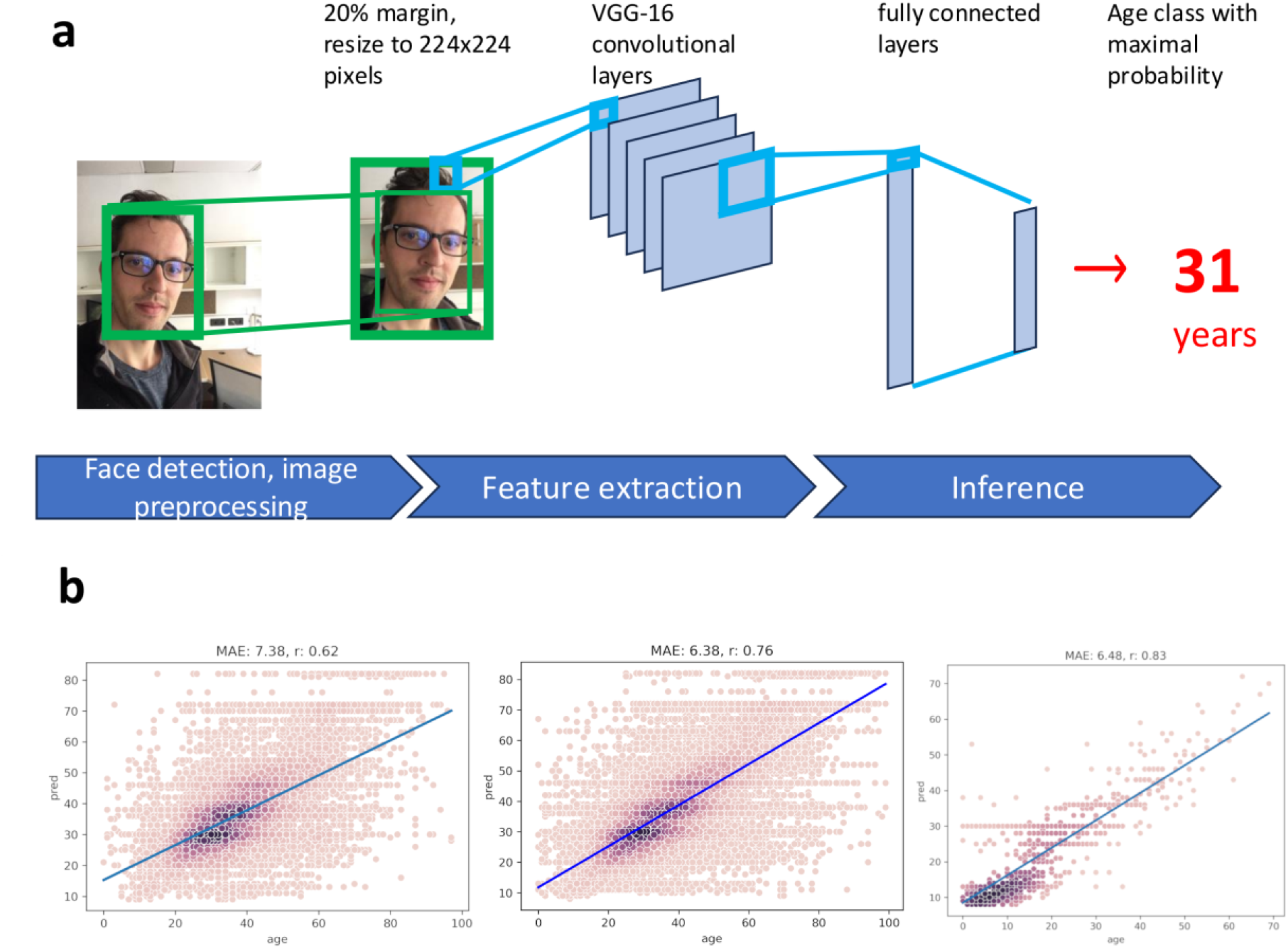
Face photo-based age prediction model trained on an automatically collected database of famous people. **a**, The pipeline of the proposed face photo-based age prediction method. It starts with face detection and image preprocessing including cropping the face, adding a 40% margin, and resizing to 224×224 pixels. Then the pipeline applies the VGG-16 convolution neural network model trained on 50% of the 442,110 cleaned images of the IMDB-WIKI database, finally, it assigns an age (in integer years between 0 and 100) associated with the maximum probability. The displayed picture shows the first author of this study. **b**, The age prediction performance of the model was measured in mean absolute error (MAE) and Pearson correlation coefficients (r) for the IMDB-WIKI test, IMDB-WIKI exploratory clean, and FG-NET datasets, respectively. Colors indicate the density of the dots (the darker the denser).

For exploratory analysis, we reduced noise by dropping the photos from the exploratory test set where the face detector algorithm detected more than one face to avoid the selection of another face than the subject. As we expected the age prediction performance of the noise-reduced (“cleaned”) exploratory set highly improved (r = 0.76 and MAE = 6.38 years) compared to the test set, however, the process dropped approximately half of the photos including wrong and right photos (Table 1, Fig. 1b). We used the cleaned exploratory set in the mortality and occupation analysis arriving to more reliable results.

### Age acceleration based on face photos predicts all-cause mortality of middle age or older famous people

We examined the relationship between the face photo-based age acceleration and all-cause mortality for famous people. The Cox proportional hazard regression on the cleaned exploratory dataset showed that one-year increase in face photo-based age acceleration is associated with a 0.8% increase in the risk of all-cause mortality (HR = 1.008, p = 1.4e-12, see Fig. 2). This effect remained significant for middle-aged and older people (age ≥ 45 years; HR = 1.01, p = 9.1e-14) but not for young people (age < 45 years; HR = 1.0007, p = 0.75). Photo-based age acceleration was associated with all-cause mortality for middle-age and older males and females, separately (HR = 1.009, p = 7.7e-10, and HR = 1.01, p = 0.0016, respectively).

**Fig. 2.**
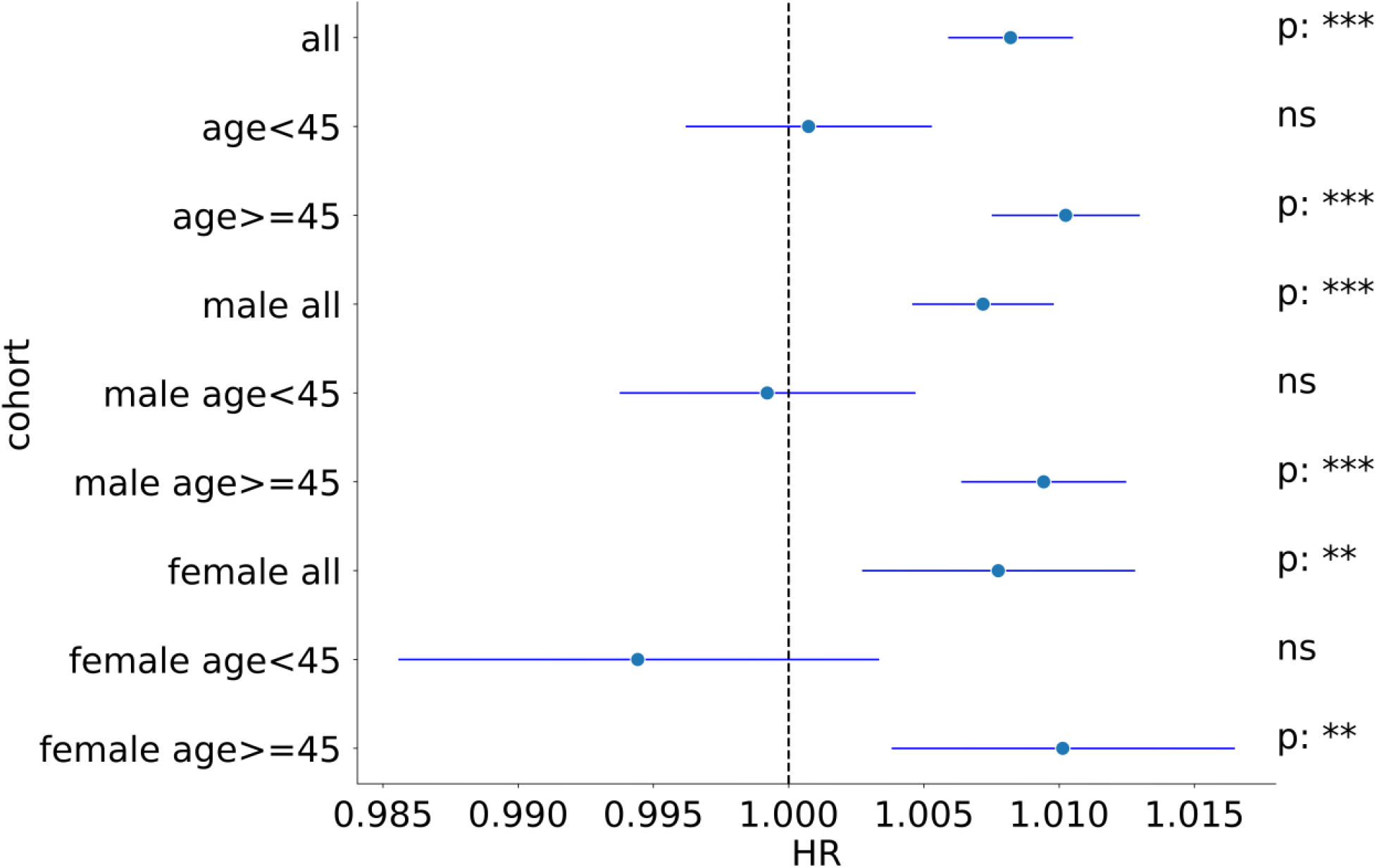
Age acceleration based on face photos predicts all-cause mortality of middle-aged and older famous individuals. Univariate (unadjusted) Cox proportional hazard regression of all-cause mortality for each subset of the cleaned exploratory set. Each row shows the hazard ratio (HR) and the 95% confidence interval for the corresponding subset. P-values are also indicated.

We also performed a survival analysis for each 15-year age interval between 30 and 90 years. In the case when more than one image was available for the same person in a given interval, we considered the image associated with the median predicted age. Individuals were considered fast agers if their median face photo-based age acceleration was greater than 5, and slow agers if it was less than -5. The survival rates of fast agers were significantly lower compared to the slow agers for the middle-aged and older age groups (45-60, and 60-75 years) but not for the young and very old age groups (30-45, and 75-90 years; see Fig. 3A). The sex-separated analysis showed similar patterns, however, surprisingly, for young males we found a significantly better survival of the fast ager group compared to the slow agers (Fig. 3BC).

**Fig. 3.**
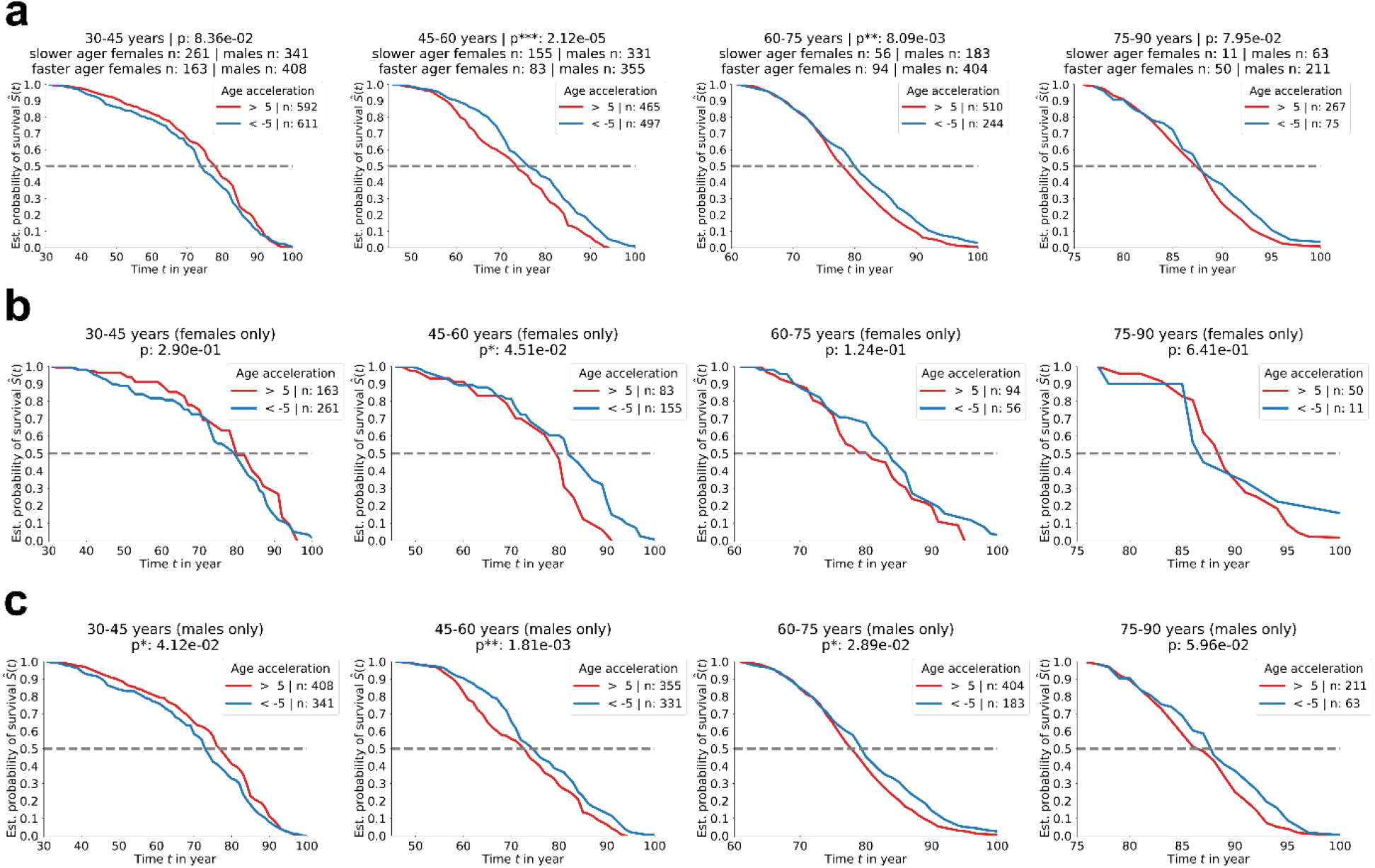
Survival analysis of fast-ager and slow-ager famous people based on their face photos. Kaplan-Meier survival analysis (all causes of death) comparing fast agers (individuals with median face photo-based age acceleration > 5), and slow agers (individuals with median face photo-based age acceleration < -5) among famous people in each 15-year age group between 30 and 90 years, for **a**, all, **b**, only females, **c**, only males. *Time t in year* indicates the actual age of the survivors.

### Athletes age the slowest based on face photos while science/education were the fastest ager occupation category among famous people

As aging can be influenced by lifestyle factors we hypothesized that aging rates may alter among occupations. To test this idea, we calculated the median photo-based age acceleration for each occupation on the cleaned exploratory dataset. We observed that the top 11 occupations with the lowest face photo-based age accelerations were some physical sports (Fig. 4). The individuals who aged the slowest were badminton players, table tennis players, and volleyball players. On the other hand, there were no sports in the top 30 occupations with the highest face photo-based age acceleration. The top three fastest ager occupations were the financier, literary critic, and historian.

**Fig. 4.**
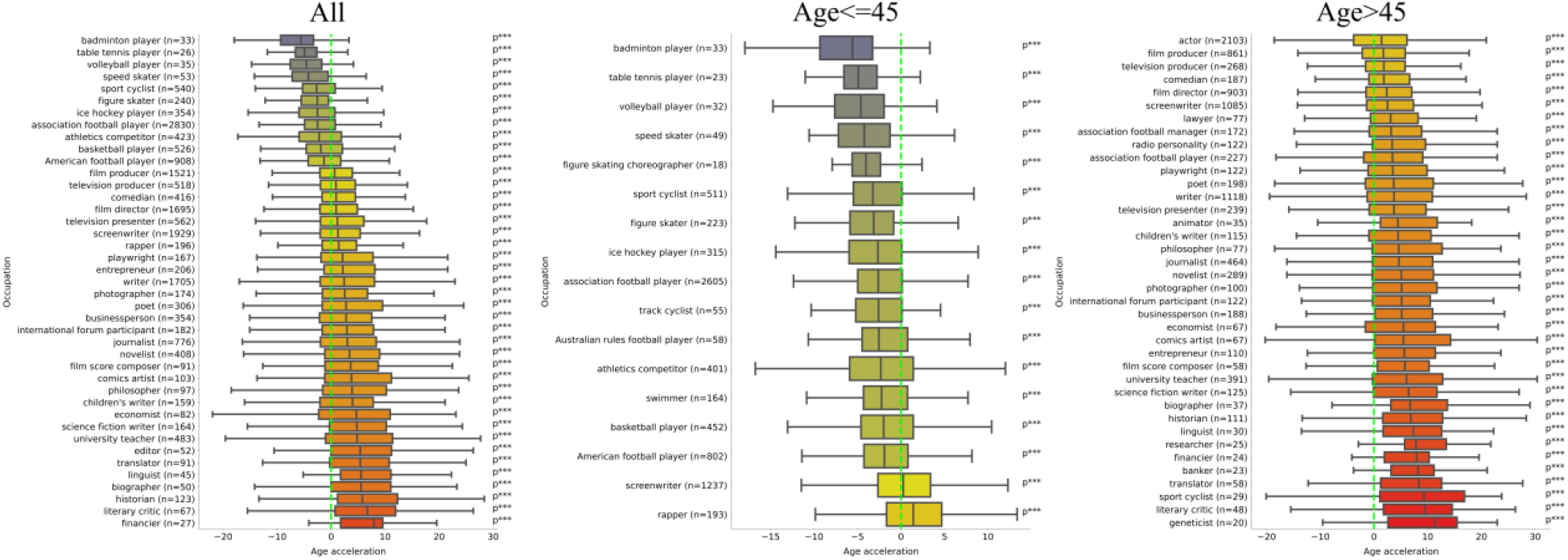
Athletes age the slowest based on face photos among famous people. **a**, Occupations where the median age acceleration based on face photos significantly differed from zero; for famous people with all age, older than 45 years, and younger than 45 years, respectively (Bonferroni correction by n = 317, 228, and 223, respectively).

We also assigned occupations to eight broader categories and found that the slowest ager occupation was again the “Sports” category and the fastest ager occupation category was the “Science/Education” (Fig. 5a). We found that for both sexes the lowest and highest age acceleration was again associated with the sports and scientific/education occupation categories, respectively (Fig. 5b). We also found that the mean age acceleration is lower for females compared to males for every examined occupation category and individual occupations.

**Fig. 5.**
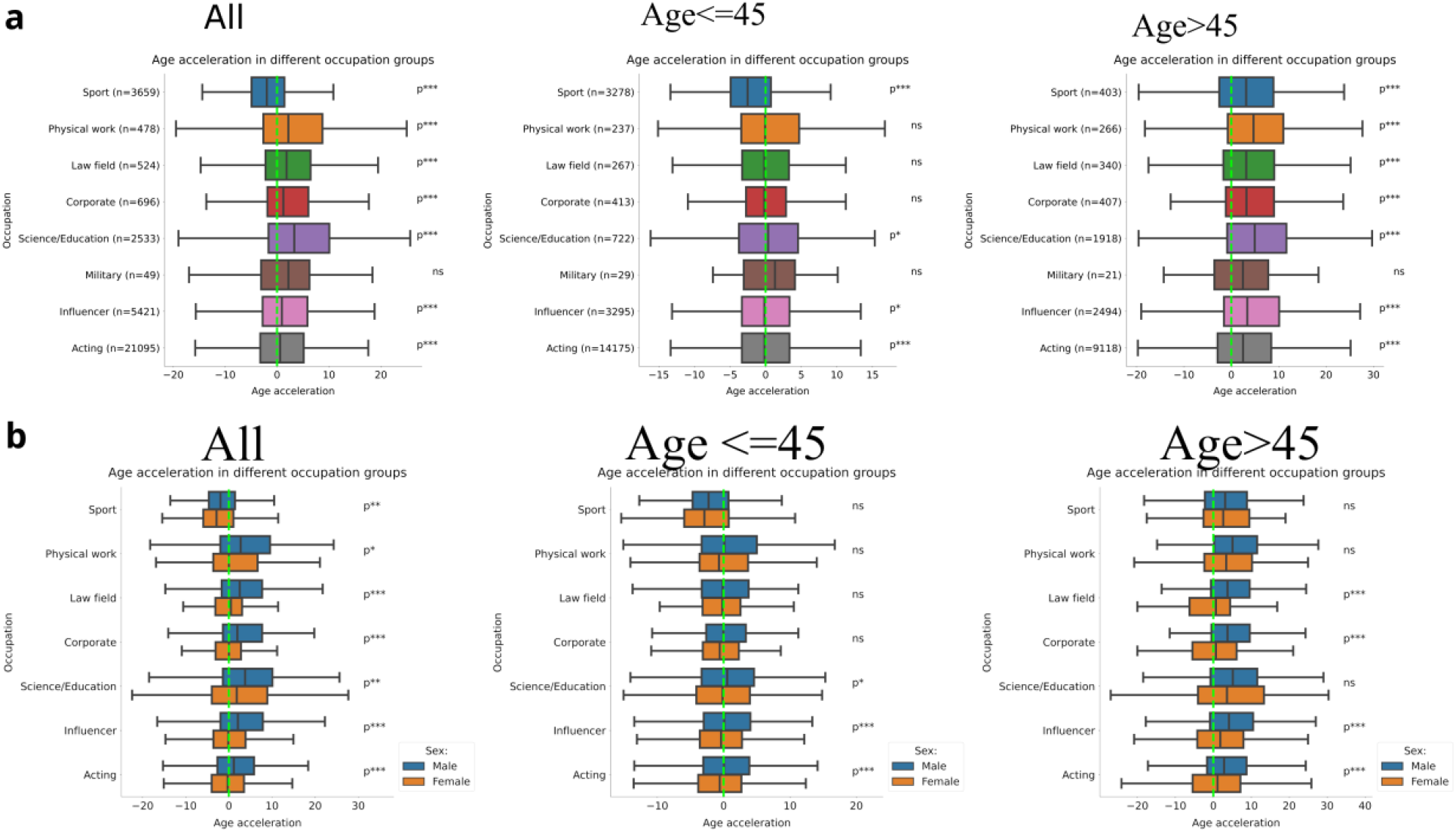
Age acceleration differences of occupation categories and sex among famous people. **a**, Age acceleration among eight broader occupation categories. The p-value indicates the significance of the deviation of the mean age acceleration for zero (two sided one-sample t-test with μ = 0). **b**, Age acceleration of the occupation categories separated by sex. The p-values of the age acceleration differences between males and females are calculated by two-sided t-tests. Only the occupations categories with p < 0.01 and n > 5 are displayed.

### Nose-mouths area was more important in the age prediction model than the eyes or the entire face

We investigated which area of the face was the most important in the age prediction of the model. We applied the Grad-CAM^17^ algorithm that can evaluate the importance of each pixel of a photo for the age prediction model. After manually checking the Grad-CAM values of some photos, we decided to examine three areas: the entire face, the eyes, and the nose-mouths area (Fig. 6a). Then we calculated the mean importance, measured by Grad-CAM values, of the areas in different age groups and datasets. The eyes area and the entire face area showed similar mean importance, while the nose-mouth area showed remarkably higher mean importance in every examined age group (Fig. 6b).

**Fig. 6.**
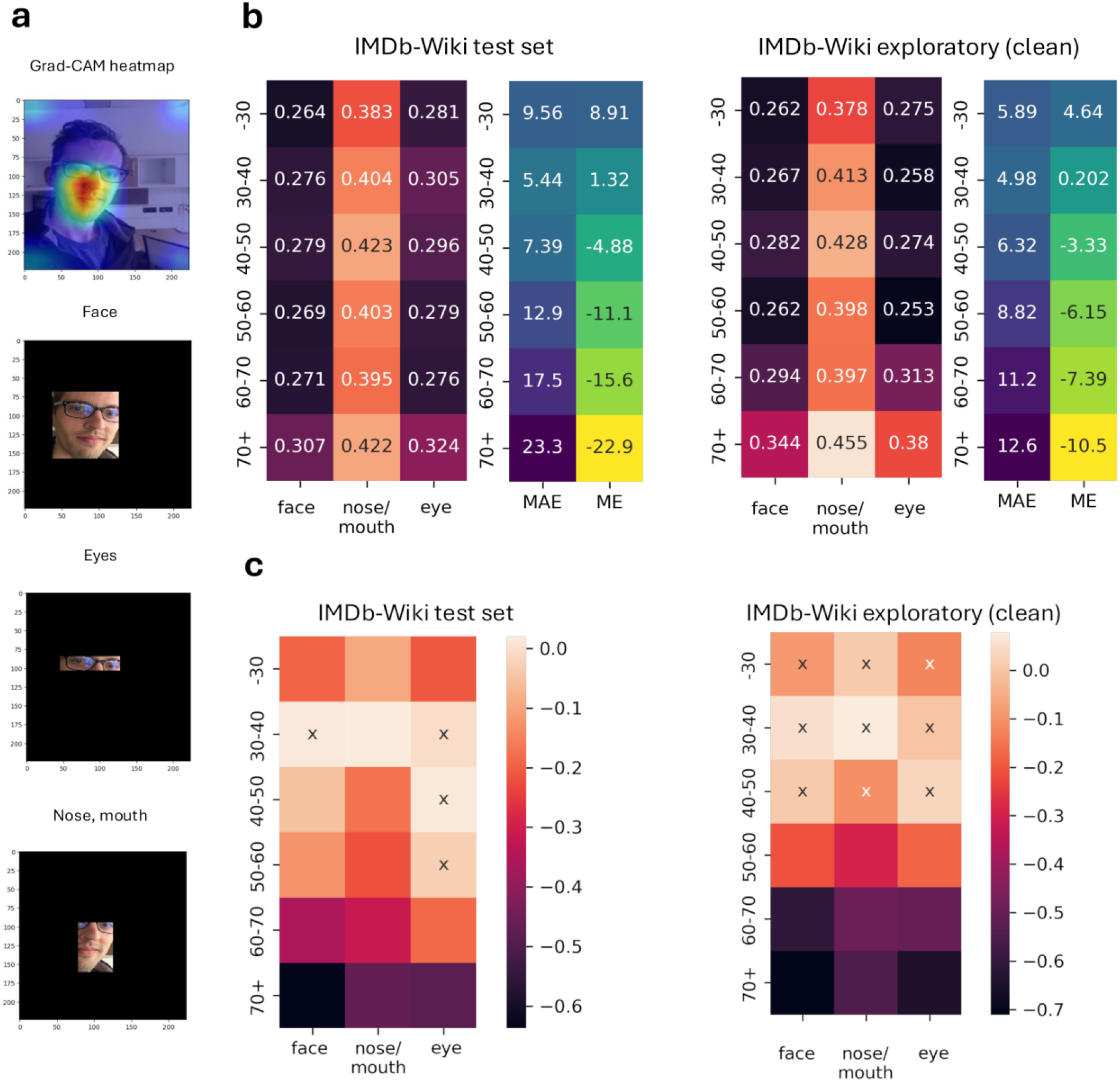
The nose-mouth area is more important in the age prediction model than the eyes or the entire face. **a**, Distribution of the Grad-CAM values (i.e., importance in the age prediction model) for an example photo and the selected facial regions. The displayed pictures show the first author of this study. **b**, Mean Grad-CAM values of facial areas on different age groups, calculated on IMDB-WIKI subsets. MAE of the age prediction is also shown for each age group. **c**, Spearman correlation coefficient between regional Grad-CAM value averages and absolute prediction errors. Nonsignificant correlations are denoted by *x*.

## DISCUSSION

For the first time, here, we showed that face photo-based age acceleration calculated by AI is predictive to all-cause mortality. These results suggest that face photo-based age can be a cost effective complementary measurement of biological age.

Interestingly, we found that face photo-based age acceleration is predictive to mortality only for middle-aged and old people (older than 45 years), and not for younger ones (below 45 years). Previous studies showed that age acceleration based on epigenetics, brain MRI, and blood tests is predictive of mortality, however, mostly old samples were analyzed and did not separate their analysis to young and old^8,18–24^. Consequently, it remained an important open question whether age acceleration in general is predictive of all-cause mortality for young people. Here, we provided a negative answer regarding the face photo-based clocks. One possible explanation of our results is that young people have more time to change their aging rates by appropriate lifestyle habits.

Our study is the first that investigated the aging rates of different occupations based on face photos. Analyzing plasma proteome aging showed that occupations requiring higher education and high autonomy, such as scientists, managers, teachers, and medical doctors, are associated with slower aging rates compared to low-paid jobs, such as cleaning workers, factory workers, and telephone operators^25^. Higher education and income were associated with accelerated epigenetic age by several epigenetic clocks^8,21,22,26^. A possible explanation of this is that people with low income and low education, which are strongly parallel characteristics, may have less opportunities to lead a healthy lifestyle and less access to healthcare benefits. Indeed, it was shown that income and education were strong predictors of mortality in different populations^27,28^. The specific cohort of the current study consists of only famous people who appeared in the Wikipedia and the IMDB database. Low-income and low education are highly underrepresented in this specific cohort. Famous people with low income are usually highly educated while those with low education have high incomes. In summary, occupations of famous people with the highest photo-based age acceleration probably would not be in the top among the general population as at least low-paid, repetitive jobs would exceed them).

Previously it was shown that regular exercise is associated with slowed epigenetic aging^2,6,29^. Analyzing Hungarian athletes, we showed that blood test-based age acceleration is inversely associated with high-volume sports activity^30^. Consistent with these results, here, we found that photo-based age acceleration of athletes is not only significantly lower than zero but athletes age the slowest among occupations.

The most accurate age prediction models were able to achieve 2-3 years MAE on special datasets, such as the mugshot images of the MORPH and the FG-NET database^31^. However, the IMDB-WIKI database is collected automatically (i.e., without human curation) from the IMDB and the Wikipedia databases, consequently, it contains highly diverse “in-the-wild” photos with substantial noise. The authors of the IMDB-WIKI database and the subsequent studies used it only for pre-training^10,12^, and not for testing. Recently, it was identified that the main source of the noise in the IMDB-WIKI database was that the relatively weak face detector method used by the IMDB-WIKI database failed to detect the right face when encountering multiple faces^14^. Consistently, the age prediction model was more accurate when we considered only the images where only one face was detected. Another source of the noise may be that many images were from movies that could have extended production times^32^. Possible photo modifications, plastic surgery, and hair color alterations are additional factors that could increase prediction error. However, these factors probably infuence only the minority of the photos. Despite the noisy dataset that we used for training and testing, our age prediction model achieved a good prediction performance on the cleaned exploratory dataset emphasizing the strong prediction power of our model. Other photo databases such as MORPH2 are less noisy and better age prediction performance can be achieved, however, they do not contain famous people and mortality or occupation data are not available. In summary, while in-the-wild photos of famous people have an inherent noise and obvious sample bias, they have the invaluable benefit of having much more available information, most importantly mortality and occupation data.

We applied the Grad-CAM algorithm to determine the importance of each pixel of a photo for the age prediction model. In the older age groups, the mean Grad-CAM values negatively correlated with the absolute prediction error. The inference quality in younger cohorts seems to be less affected by the low facial Grad-CAM values – possible explanations to this could be the bias of the training dataset towards people in their thirties (Fig. 6c).

Another possibility to approach face photo-based biological age is perceived age, which is defined as how old a person looks according to external human evaluators^33^. Cox proportional hazard analyses of multiple studies showed that sex and chronological age adjusted perceived age predicted mortality of 70+ year old people^34–36^. Compared to perceived age, our AI-based approach may provide a much higher scale, more robust, and reproducible way for the assesment of face-photo based biological age.

Overall we developed a face photo-based aging clock model that can be a complementary measurement of biological age and provides a cost-effective and fast tool for future personalized medicine, as well as aging and rejuvenation studies.

## METHODS

### Face image datasets

The IMDB-WIKI is a large-scale dataset of face images, consisting of 523,051 images from celebrities (460,723 face images of 20,284 individuals from IMDB and 62,328 from Wikipedia) labeled with the corresponding ages (https://data.vision.ee.ethz.ch/cvl/rrothe/imdb-wiki/)^10^. The dataset also contains the location of the face on the image as well as detector scores of the detected faces corresponding to the two highest values. In our analysis, we used a version of the database where each image was cropped around the highest-scored face, padded with a 40% margin around it. In the cases when the face covered most of the image it was padded with the last pixel at the border. It ensures that the face remains in the middle of the image. We dropped zero-pixel, and corrupted images (where checksum failures were detected in the image file) as well as those that were labeled older than 100 or younger than 0 years, and images where the face detection score was *Inf* (i.e., no face detected). After these preprocessing steps, we split the remaining 442,110 images into two subsets in a proportion of 50-50%. The first half was used for model development. 70% of images from the first half were used for training, 15% for validation, and 15% for testing the models. The second half was used as an additional testing dataset for explorations. The splits were made person-wise i.e. there was no person whose face images were included in more than one subset. For exploratory analysis, we dropped the photos from the exploratory test set where the face detector algorithm detected more than one face to avoid the selection of another face than the subject. We refer this as “cleaned exploratory dataset”.

The models were also evaluated on the FG-NET dataset containing 1,002 curated mugshot images of 82 non-celebrity individuals with age between 0 and 69 years (https://yanweifu.github.io/FG_NET_data/).

### Face photo-based age prediction model development

Age prediction was formulated as a classification problem using age groups for every year between 0 to 100. Here, we used the VGG-16 architecture for the classification task, which was initialized with the weights pre-trained on the ImageNet^37^. Input images (cropped faces with 40% margins) were resized to 224×224 pixels and then normalized by the means and standard deviations of the ImageNet RGB channel. We used a dropout rate of 0.7 and replaced the last layer with 4096 fully connected neurons. For each experiment, we used an initial learning rate of 0.0001, which is multiplied by 0.5 if the validation loss stopped decreasing. Adam optimiser was used, and the training process was early stopped when the model performance did not improve in the last 50 epochs.

### Evaluation of the model performance

The best-performing model on the validation set was also tested on the testing dataset, the cleaned exploratory dataset, and the FG-NET dataset. We evaluated the models by calculating the mean absolute error (MAE, in years) and the Pearson correlation coefficients (r) between the age predictions and chronological ages (Table 1).

### Age acceleration

We computed age acceleration for all face photos of the cleaned exploratory dataset as follows: we fit a regression line to the predicted age as a function of chronological age, then for each face photo, the age acceleration was calculated as the difference of the predicted age and the value of the regression line. Age acceleration is independent of age by definition. If the age acceleration is negative we assume slowed aging, if it is positive, we assume accelerated aging for the given person.

### Mortality and occupation data

We downloaded the Crossverified Database of Notable People^18^, which contained information from all human-titled Wikipedia pages, and found the Wikidata IDs of the individuals who appeared in our exploratory dataset. Wikidata is a large public database that serves the entire Wikipedia. Using the Wikidata IDs, we collected lifestyle data (date of death, cause of death and occupation) from Wikidata.

### Cox proportional hazard regression analysis

In Cox’s proportional hazard model, the log hazard is a linear function of the covariates *x*_*i*_and a population-level baseline hazard *h*_0_:

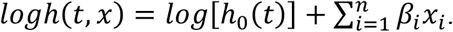

In our analysis, the event was the death of an individual associated with a given photo. We considered every photo as a different individual. The predictor variables were age, predicted age, and sex. We conducted separate analyses for different subsets of the exploratory dataset: all photos, below 45 years, above 45 years, males, and females. The p-values were calculated by Wald tests.

### Kaplan-Meier survival analysis

Kaplan-Meier survival analysis examines the occurrence of an event in a population. Here, the event is the death of a person. The Kaplan-Meier estimator curve is a non-parametric method for approximating the survival function. One of the main advantages of the estimator is that it can handle censored data. The probability of survival after *t* is

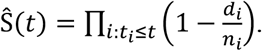

Where Ŝ approximates the survival function as a function of *t*, the t_i_ denotes a time point when at least one event occurred, d_i_ is the number of events that occurred at t_i_, and n_i_ is the number of surviving samples. We performed a Kaplan-Meier survival analysis for each 15-year age interval between 30 and 90 years. In the cases when more than one photo was available for the same person in a given interval, we considered the photo associated with the median predicted age. Individuals were considered fast agers if their median face photo-based age acceleration was greater than 5, and slow agers if it was less than -5. We compared the survival rates of the two groups by using a logrank test which is a nonparametric hypothesis test that compares the survival distribution of two statistical samples.

### Model explanation

To get an insight into the mechanisms of the models, we used the Grad-CAM algorithm^17^. We examined its output when applied to the final layer of the models. The output can be represented as a matrix of the same size as the input image with values from 0 to 1 representing the contribution of each pixel to the prediction. In order to locate the facial area the model pays the most attention to, we bounded three areas on each image of the IMDB-WIKI test, and the cleaned exploratory datasets where the Grad-CAM weights showed notable concentration. These areas were the entire face, the eyes, and the nose-mouth region including the nasolabial fold. For each area, Grad-CAM output matrices were cropped according to the corresponding bounding boxes obtained by the Google MediaPipe^38^ built-in face landmarkers, then the averages of the remaining values were calculated.

### Statistics

Mean Absolute Error (MAE) is a widely used metric for comparing models used for age estimation.

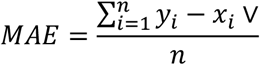

Where n is the number of individuals; x_i is the chronological age and y_i is the predicted age of the ith individual.

Pearson correlation coefficient (r) is also a commonly used metric in the evaluation of age prediction models. The r is a numerical value between -1 and 1 that measures the linear correlation between two variables.

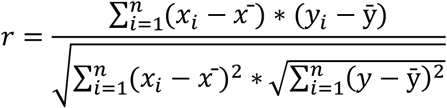

Where n is the number of individuals; x_i_ is the chronological age and y_i_ is the predicted age of the ith individual; 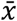 and ȳ are the averages.

If p values were indicated by an asterisk, we used the notations as follows: ns, p > 0.05; *, 0.01 < p ≤ 0.05; **, 0.001 < p ≤ 0.01; ***, p ≤ 0.001, ****, p ≤ 0.0001. Density coloring of the scatter plots was performed using the values of the probability density function obtained by Gaussian kernel density estimation. Computational experiments were carried out using PyTorch 1.13.1 and Torchvision 0.14.1.

## DATA AND CODE AVAILABILITY

The face-photo-based age prediction model is available for academic research purposes at https://photoage.sztaki.hu/. Codes are available on Zenodo (https://doi.org/10.5281/zenodo.14031057).

## ACKNOWLEDGEMENTS

The project was supported by the European Union project RRF-2.3.1-21-2022-00004 within the framework of the Artificial Intelligence National Laboratory, and the National Research, Development and Innovation Office – NKFIH, FK-146113.

## AUTHOR CONTRIBUTIONS

CK conceived and supervised the study. KB developed the photo age prediction model, evaluated the performance, and explained the model. IF performed the mortality and occupation analysis. All authors wrote the manuscript.

